# Modular fluidic automation platform with integrated thermal control for multi-step molecular imaging workflows

**DOI:** 10.64898/2026.04.17.713973

**Authors:** Tirtha Das Banerjee, Joshua Raine, Ajay S. Mathuru, Antónia Monteiro

## Abstract

Automation of multi-step mRNA imaging protocols increases reproducibility and throughput in spatial biology, as many workflows require repeated buffer exchanges, precise timing, and controlled reaction conditions. Commercial automation platforms can be expensive, proprietary, and difficult to customise, limiting their use in most laboratories. Here, we present two open-source robots for the Rapid Amplified Multiplexed Fluorescent In-Situ Hybridization (RAM-FISH) workflow based on programmable delivery of fluids and integrated thermal control with no dedicated bubble trap requirement. The first robot is designed to perform the steps necessary for signal localization (Multiplexer), and the second performs signal removal (RemBot). Both robots function without manual supervision and conduct precise, repeatable buffer exchanges, temperature regulation, and timed reactions. Both can operate on free-floating and gel-embedded tissues and can be assembled using widely available components. The robots support iterative imaging workflows, enabling detection of multiple genes across sequential hybridization rounds within the same sample. By providing customizable and accessible robots, we lower the technical know-how barriers that need to be overcome to perform complex spatial imaging experiments and enable scalable, hands-free execution of multi-step multiplex-FISH.

## Introduction

Automated fluid handling has become increasingly important for modern molecular imaging and spatial biology workflows, which often require multiple cycles of reagent exchange, precise incubation timing, and controlled reaction environments. Techniques such as multiplexed-Fluorescence In Situ Hybridization (multiplexed-FISH), immunostaining, and sequential labeling involve repetitive and time-sensitive steps that are labor-intensive and prone to variability when performed manually. These challenges limit experimental throughput, increase user fatigue, and introduce inconsistencies that can affect signal quality and reproducibility.

Commercial automation platforms such as MERSCOPE and Xenium are highly effective for multiplex-FISH assays, however, such systems are often expensive, proprietary, and optimized for specific applications or consumable formats (Chen et al., 2015; Janesick et al., 2023). Such systems frequently lack flexibility for customising experimental workflows, integrating diverse sample types, or compatibility with standard microscopes and laboratory infrastructure. As a result, many research laboratories continue to rely on manual sample processing for performing such multi-step protocols, particularly in academic settings where experimental conditions are frequently updated.

Recent advances in open-source electronics, microcontrollers, and rapid prototyping have enabled the development of customizable robots that are accessible and adaptable (Deng and Beliveau, 2022; Pearce, 2012; Pearce, 2014). Simple robots with programmable fluid delivery, temperature control, and user-defined timing can provide a practical alternative to commercial platforms, allowing researchers to tailor automation to specific experimental needs while reducing cost and technical barriers. Currently available open-source systems (Deng and Beliveau, 2022; Moffitt and Zhuang, 2016), however, have not been applied to thick 3D tissues and lack thermal control critical for multiplexed-FISH reactions such as HCR3.0 (Choi et al., 2018; Gandin et al., 2025).

With cost, flexibility, and diversity of samples as our motivating factors we developed two open-source robots that can conduct the multi-step mRNA detection workflow, RAM-FISH, with minimal user involvement (Banerjee et al., 2024). Here, we provide detailed instructions for assembling these robots (Multiplexer and RemBot) using widely available components and a 3D printer. The robots employ microcontroller-driven architectures that enable precise timing, repeatable reagent delivery, and stable temperature regulation of free-floating or gel-immobilized samples within a standard 35 mm dish. The fluidic design enables quick replacement of tubing (in the event of clogging due to salt crystallization) and eliminates the requirement for dedicated bubble-trapping components, simplifying system maintenance and operation. The functionality and usability of these robots have been tested extensively with users across a range of expertise levels, indicating ease of adoption and adaptability in typical laboratory settings. The spatial localization of dozens of mRNAs and microRNAs per sample in different species (*Bicyclus anynana* larval wings and 14 dpf *Danio rerio* larval brains) suggests broad usability (Banerjee and Monteiro, 2025; Goel et al., 2026; Prakash et al., 2024; Raine et al., 2025; Tian et al., 2024) over 8-10 cycles of probe-RNA hybridization, imaging, and signal removal (Banerjee et al., 2024). Overall, these robotic aides, improved reproducibility, reduced hands-on time of researchers, and facilitated species-independent spatial imaging protocols.

### Workflow Overview

The complete system includes:

1. <u>Multiplexer</u>: A robot for performing multiple rounds of FISH reactions with controlled temperature in a dual dish setup.
2. <u>RemBot</u>: A robot dedicated to removing fluorescent signals from previously probed samples at a controlled temperature.
3. <u>Confocal microscope</u>: For imaging (commercial).

The workflow of a FISH experiment is illustrated in **Figure 1**.

**Figure 1.**
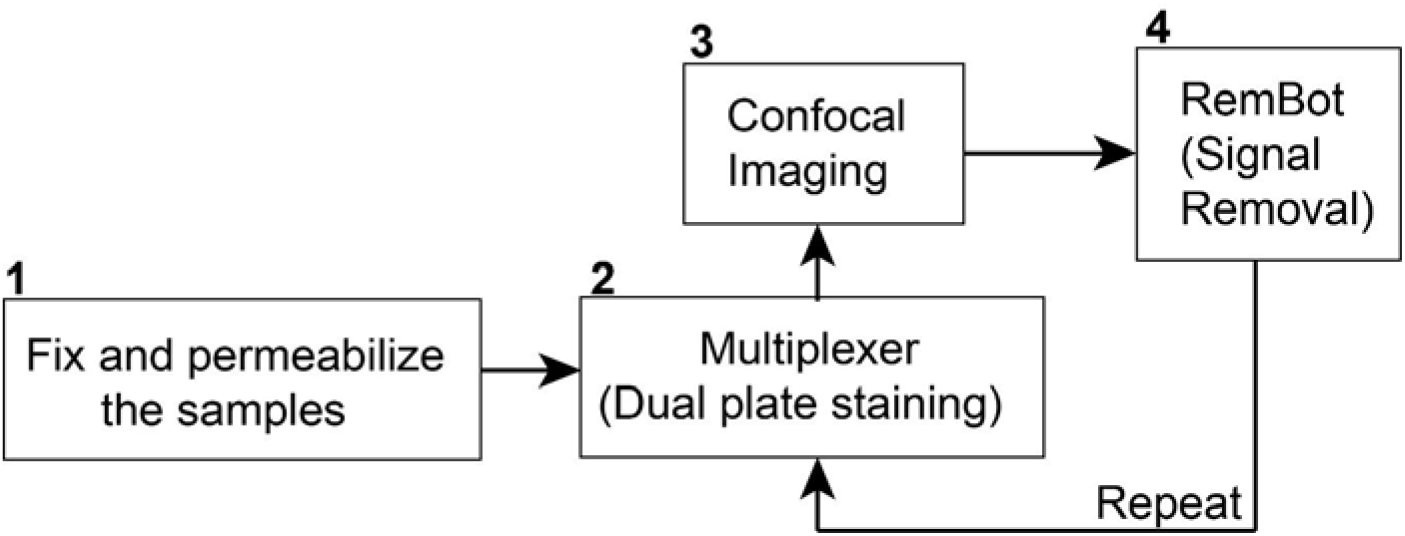
Simplified workflow of an automated RAM-FISH experiment.

### Hardware Design

The core hardware in the two robot systems includes:

#### Fluidic module

This component controls the flow of fluids into the reaction chamber using peristaltic pumps and a microcontroller. It is used in both the Multiplexer and RemBot robots.

#### Thermal module

This component includes two resistive heaters with thermistors at the bottom of the two reaction plates. It is used in both the Multiplexer and RemBot robots.

#### Mechanical components

The systems involve probe module plates for easy buffer loading and washing on top of the main body. The front panel is foldable and connected via dampening hinges that allow loading and unloading of the confocal dishes containing the samples. The confocal dishes are magnetically locked to ensure that they stay stable during the staining, imaging, and washing cycles.

#### Display

Displays the reaction steps, time, and temperature. The buttons next to the display are used for programming different reaction conditions.

### Assembly

#### Tools needed

Mini Screwdriver set: Xiaomi Electric Precision Screwdriver

Power screwdriver: BLACK+DECKER 4.8V Cordless NiCad Screwdriver With 15 Piece Bits Set KC4815

Power drill: BLACK+DECKER 8V MAX* Cordless Drill + 43 pc. Home Decor Project Kit (BDCD8HDPK)

Hot glue gun: INGCO Corded High Temperature Hot Glue Gun GG148

Measuring scale: Generic

Wire cutter/stripper: Generic

Soldering iron: INGCO electric soldering iron with solder feeder SI016732

#### Consumables needed

Hot glue refills (11×150 mm): 10 units.

Lead-free solder (Sn 99.3%, Cu 0.7%): 500 gms.

Thermal paste (Arctic MX-4): 4 gms.

Self-tapping Screws (stainless steel, assorted): 4.2×13mm, 4.2×19mm, 4.2×25mm, and 4.2×32mm.

35mm confocal dish (SPL Cat No.: 101350).

Fine nylon mesh (0.05 mm).

Double sided tape (high performance).

Below, we describe the parts, CAD designs, circuit diagram, and assembly steps necessary for building the two robots.

#### Multiplexer

The parts and instruments required for the assembly of this robot are described in **Table 1**. CAD designs for the 3D parts are in **Table 2**. Dimensions of the 3D parts are in **Figure 2**, and a circuit diagram is in **Figure 3**. The Multiplexer robot is designed for staining samples in a dual dish setup. An assembled system is shown in **Figure 4**.

**Table 1:** Parts required for the Multiplexer automation system.

| Sl. No. | Part | Configuration/Part number | Detail | Number of units/quantity |
| --- | --- | --- | --- | --- |
| 1. | Microcontroller (Microchip ATmega 2560) | Arduino Mega 2560 | Controls the pumps and communicates to the secondary microcontroller (to control the temperature) | 1 unit |
| 2. | Screw terminal adapter for Mega 2560 | Generic | Screw terminal adapter for the ATmega 2560 microcontroller allows solder-free connection of the wires to the microcontroller's IOs. | 1 unit |
| 3. | Secondary microcontroller (Microchip ATmega328p) | Arduino Nano 328p (USB-C connector) | Controls the heating plates | 1 unit |
| 4. | Screw terminal adapter for Nano 328p | Generic | Screw terminal adapter for the ATmega328p microcontroller allows solder-free connection of the wires with the microcontroller IO. | 1 unit |
| 3. | 12V DC peristaltic pumps (7W) | DC 12V, Flow rate:0-100 ml/min, Tube size: 2mm ID x 4mm OD | Controls the inlet and outlet liquid dispersion and removal in the confocal dishes. | 18 units |
| 4. | Pump driver board | L298N | Dual H-bridge motor drive module. Controls the peristaltic pumps' speed and direction. | 9 units |
| 5. | 4-bit button module | Generic | Buttons to run a specific program. | 1 unit |
| 6. | 2 channel 5v relay | Maximum contact: AC 250V 10A or DC 30V 10A. | Controls the dual heater plate from the secondary microcontroller. | 1 unit |
| 7. | 1 channel 5v relay | Maximum contact: AC 250V 10A or DC 30V 10A. | Controls when to heat the thermal plates. | 2 units |
| 8. | LCD2004 Module I2C Interface 5V | Green LCD2004+ converter Module PCF8574 | Displays the temperatures. | 1 unit |
| 9. | 3.5 inch 320x480 IPS Capacitive Touch Screen | SPI Serial ST7796U Driver | Main display that shows the steps (with animations). | 1 unit |
| 10. | Colored wires male-female (50cms) | 22 AWG | For connection. | 50 wires |
| 11. | Colored wires male-female (30cms) | 22 AWG | For connection. | 80 wires |
| 12. | Solid core wires (red, black, green, yellow, blue) | 22 AWG | For connections that require higher current (such as during heating and cooling) | 20 meters |
| 13. | Translucent high performance Silicone Tubing, | (ID - 1mm; OD - 2mm) | Main tubing that connects the source, chamber, and sink. | 5 meters |
| 14. | Translucent high performance silicone tubing, | (ID - 1.6mm; OD - 3mm) | Tubing to join the 1 mm tubes. | 1 meter |
| 15. | Translucent high performance silicone tubing, | (ID - 2 mm; OD - 3mm) | Tubing for microcomb outlet. | 10 cms |
| 16. | Y-connector | (3.5mm) | To connect the tubes | 10 units |
| 17. | Heating element | (21x35 mm, 12V, 25W): Resistive heaters | For heating the plates. | 2 units |
| 18. | Thermistor+10k ohm resistor | - | Measures temperature of the heating elements. | 3 units |
| 19. | Damping hinges | 36x28mm: 2 units | Used to open and close the front panel during loading and unloading of the confocal dishes. | 2 units |
| 20. | Self-tapping screws | Stainless steel (assorted): 4.2x13mm, 4.2x19mm, 4.2x25mm, and 4.2x32mm | For connecting the peristaltic pumps and the 3D printed parts. | 50 units |
| 21. | USB cables | USB-A to USB-C, and USB-A to USB-B | USB-A to USB-B, and USB-A to USB-C | 2 unit |
| 22. | Power supply | 8-10A, 12V | To provide power to the system | 2 units |
| 23. | DC-DC power converter | Input: 3.0-40V to Output: 1.5-35V | To convert 12 V to 5V for powering the microcontrollers, display, and thermistors. | 1 unit |
| 24. | Power switch | 250V, 6A | To connect the 12V live wire from the power supply to the 12V common for the system. | 1 unit |
| 25. | PLA filament (black-1 spool and grey-2) | BambuLab PLA filament with RFID | For 3D printing. | 3 kgs |
|  | spools) | spools. |  |  |
| 26. | 3D printer | BambuLab H2D Pro 3D printer + AMS 2 Pro | 3D printer for printing parts of the system | 1 unit |

**Table 2.** CAD parts for the Multiplexer system (CAD files available at: https://github.com/tdblab/RAMFISH/tree/main/Multiplexer_STL).

| Sl. No. | Part | Description |
| --- | --- | --- |
| 1. | Main body | The main body of the system that holds the electronics and the pumps. |
| 2. | Front panel | Holds the display and buttons to control the system. |
| 3. | Top panel | Holds the probe modules with the buffer at the top of the system. |
| 4. | Back panel | Back plate of the system. |
| 5. | Chamber body | Main chamber that holds the confocal plate, confocal dishes, and the buffer exchanges. |
| 6. | Chamber body bottom plate | Bottom plate of the chamber body. |
| 7. | Confocal plate (136x96 mm; 125x85 mm; 125x80 mm) | Confocal microscope specific plate (designed differently for different microscope models) |
| 8. | Magnetic clip | Magnetic clip to hold the confocal dishes in the plate. |
| 9. | Microcomb adapter | Adapter to allow inlet and outlet to the confocal dishes |
| 10. | Microcomb plate | Plate to allow inlet and outlet to the confocal dishes (dual format) |
| 11. | Microcomb holder (free floating) | Dish holder for reactions that are carried out in free floating medium. |
| 12. | Microcomb holder (gel based) | Dish holder for reactions that are carried out in acrylamide gel. |
| 13. | Probe module plate 1 (Buffer plate) | Plate to hold the buffers. |
| 14. | Probe module plate 2 | Top lid of the buffer plate connected to the tubes. |
| 15. | Bottom cylinder support | Bottom cylindrical legs of the system. |

**Figure 2:**
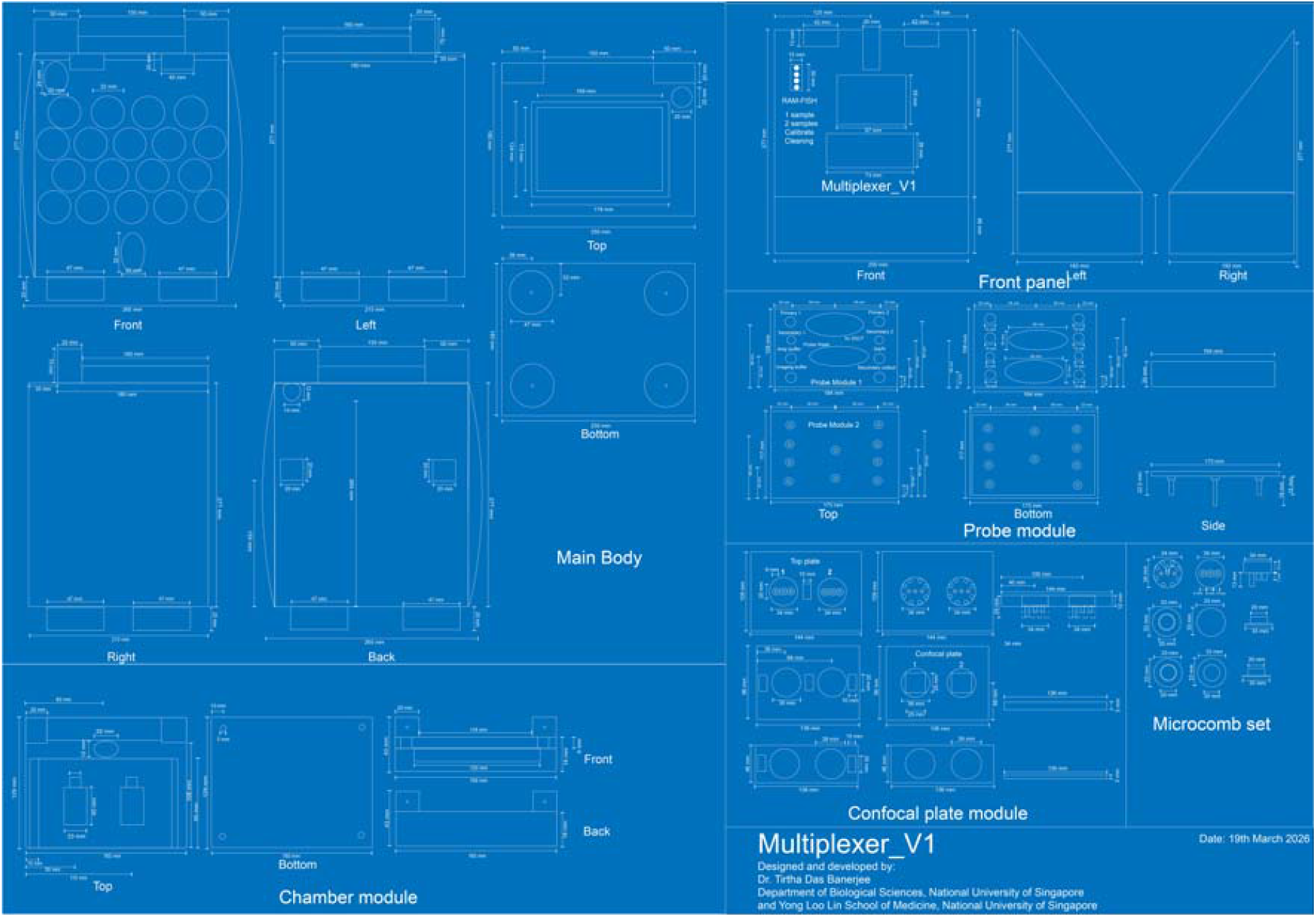
Architecture of the outer hardware of the Multiplexer robot. The components include the main body, front panel, probe module, confocal plate module, microcomb set, and chamber module.

**Figure 3:**
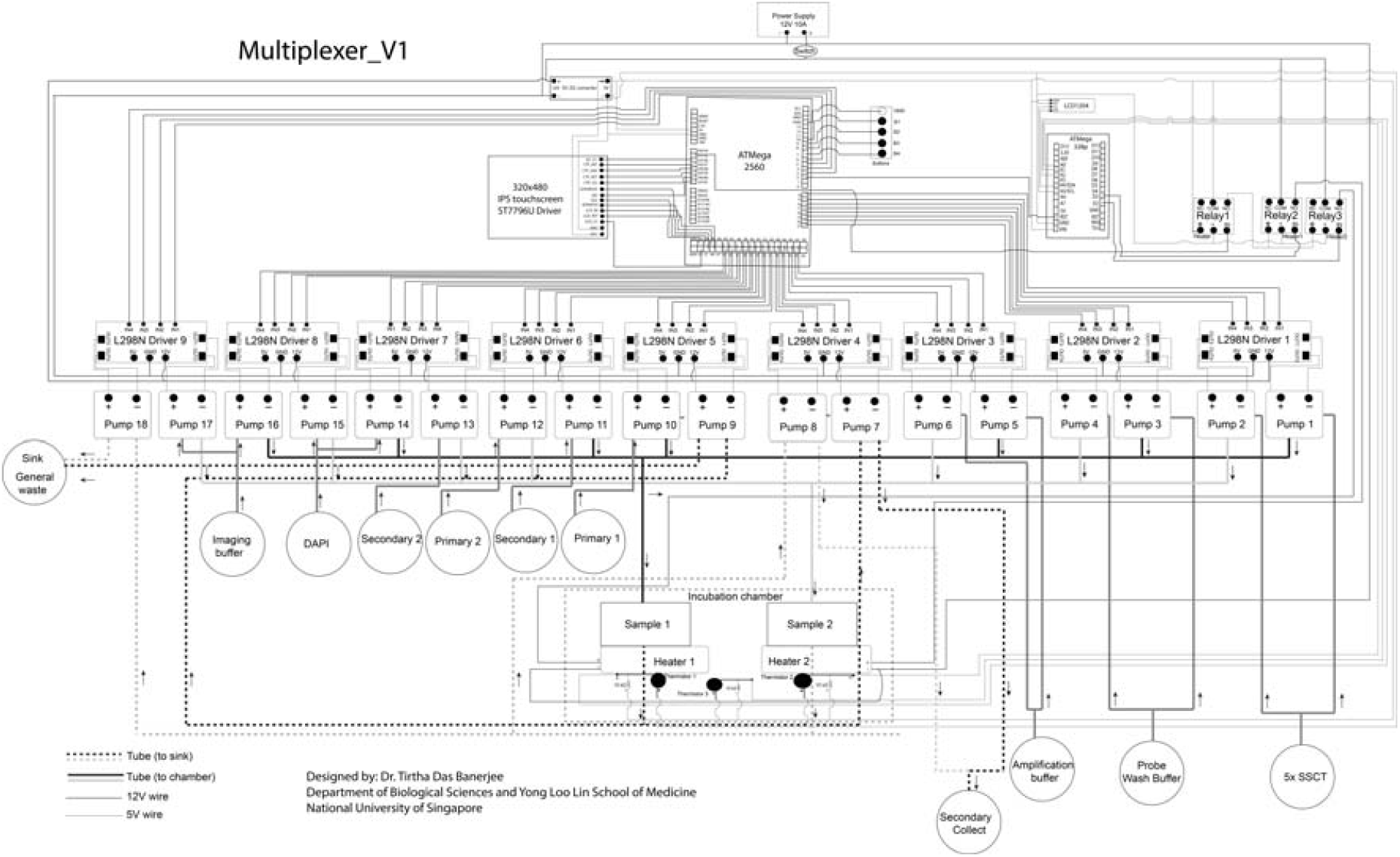
Circuit diagram of the Multiplexer robot. The architecture integrates two microcontrollers, multiple pumps, sensors, and heating plates into an integrated system for carrying out the reactions in dual dish setup.

**Figure 4:**
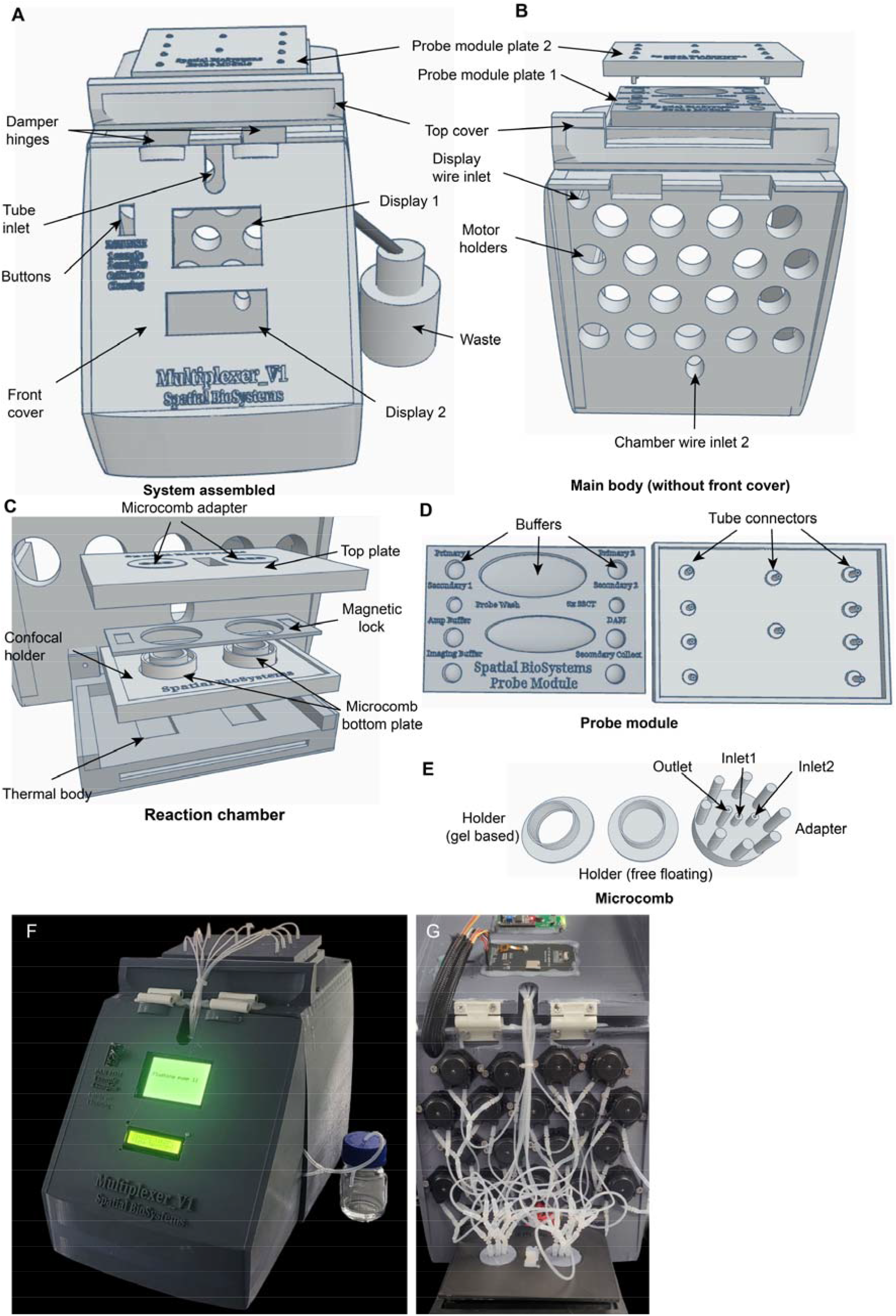
Main parts and design of the Multiplexer robot. **(A)** The main body of the robot. (**B**) Main body without the front cover showing the pump motor holders and the wire inlets. (**C**) Reaction chamber showing the microcomb top plate, magnetic lock, reaction surfaces, and the bottom plate. The thermal body holds the heating and cooling pads. (**D**) Probe module showing the plate to load the reaction buffers and the tube connectors from the top plate. (**E**) The microcomb module showing the bottom plate and top comb. An assembled Multiplexer-V1 robot. **(F)** With front panel closed and (**G**) with front panel open.

##### Step-by-step instruction for the Multiplexer robot assembly

Download all the STL files from the GitHub link (https://github.com/tdblab/RAMFISH/tree/main/Multiplexer_STL) provided in **Table 1** and 3D print them. For the assembly of all the STL files and the components in **Table 2** follow the wiring architecture in **Figure 3** and the step-by-step instructions as mentioned below:

#### Main body

1. Solder wires to the peristaltic pumps and attach the pumps using screws to the main body.
2. Attach the pump wires to the L298N driver pins. Keep a note of the orientation of the connections. Two pumps can be connected to a single driver.
3. Connect the 12 V of the L298N drivers to a common 12V and the GND of the drivers to a common GND. The 12V common is connected to one side of the power switch; the other connection is to the power supply unit.
4. Connect the data pins from the L298N controllers to the IO pins of the Mega 2560 board (Pins: 3-6 and 14-44).
5. Connect the 12V common and GND to the input of the DC-DC voltage converter, adjust the variable resistor to match 5V using a multimeter as output (Note: Do this before connecting the 5V to the microcontrollers). Connect the output 5V and GND to the common 5V and GND.
6. Install the 2-channel and 1-channel 5V relays using double sided tape in the main body and connect the 5V and GND from the common 5V and GND from the DC-DC converter. Connect the IN of the 1-channel 5V relay to pin A1 of the Mega 2560 board.

#### Front panel

7 Install the button module, IPS, and LCD screen with attached female to male wire to the front panel using hot glue.
8 Connect the 5V and GND to the VCCs and GNDs of the microcontrollers, displays, and thermistors.
9 Connect the data pins of the IPS display to the IO of the Mega 2560 board.
10 Connect the data pins of the LCD display to the I2C pins (A4 and A5) of the Nano 328p board.

#### Connecting the main body and the front panel

11 Attach the main body to the front panel using the damping hinge. The hinges can be connected using the self-tapping screws (4.2×13mm).
12 Attach the bottom cylindrical supports to the main body using the self-tapping screws (4.2×19mm).

#### Chamber

13 Attach the resistive heaters and the thermistors+10kΩ resistors assembly in the two slots provided in the chamber body.
14 Attach the chamber body bottom plate to the chamber body using self-tapping screws (4.2×13mm). Make sure the wires are accessible for later connection.
15 Attach the chamber body to the main body using self-tapping screws (4.2×32mm). Make sure the chamber body is horizontally aligned.
16 Connect 12V to the COM of the 2-channel 5V relays and NO channels to the two heaters 12V. Connect the GND from the heaters to the common GND. The IN pins from the relays are connected to the D2 and D3 pins, and the thermistor data pins to the A0 and A2 pins of the Nano 328p board.

#### Connecting the top, back, and main body

17 Connect the top panel to the main body using self-tapping screws (4.2×32mm) and the back panel to the main body using self-tapping screws (4.2×19mm). The USB cables are attached to both the Mega 2560 and Nano 328p boards before attaching the back panel.

#### Tubing assembly

18 Cut the tubes with 1.6mm ID into 1cm pieces and attach them to the outlets of the probe module plate 2. Connect the 1mm ID tubes to the 1.6mm tubes and join them to the inlets of the peristaltic pumps. Use Y-connectors with the 1.6mm ID tubes for the buffers that are shared with two pumps.
19 Connect the outlets from the pumps to the inlets of the microcomb adapters using 1mm ID tubes, 1.6mm ID tubes, and the Y-connectors.
20 Connect the outlets from the microcomb adapters to the outlet pumps (17 and 18) using 1mm ID tubes, 1.6mm ID tubes, and the Y-connectors to a 50 ml glass waste bottle.

#### Microcomb plate assembly

21 Attach fine nylon mesh (0.05 mm) to the output end of the microcomb using the 2 mm ID silicone tube (1mm height).
22 Attach the microcomb to the microcomb plate using hot glue.
23 Connect the inlets and outlets of the tubes to the microcomb inlets and outlets.
24 Add the confocal plate to the top of the chamber and install the microcomb plate.

##### Uploading the control software

Download the codes for running single and dual samples for the Multiplexer robot from the GitHub link: https://github.com/tdblab/RAMFISH/tree/main/Multiplexer_code (multiplexer.ino and multiplexer_thermals.ino). Download and install Arduino IDE from the official site (https://support.arduino.cc/hc/en-us/articles/360019833020-Download-and-install-Arduino-IDE) in your computer. Install the necessary libraries (LCDWIKI_GUI.h, LCDWIKI_SPI.h, LiquidCrystal_I2C.h). Connect the USB-A end of the USB cables to one of the PCs USB ports and the USB-C/USB-B ends to the boards. First, select the Mega 2560 board and upload the multiplexer.ino code. Next, select the Nano 328p board and upload the multiplexer_thermals.ino code.

#### RemBot

The parts and instruments required for the assembly of this robot are described in **Table 3**. CAD designs for the 3D parts are in **Table 4**. Dimensions of the 3D parts are in **Figure 5**, and a circuit diagram in **Figure 6**. The Multiplexer robot is designed for staining samples in a dual dish setup. An assembled system is shown in **Figure 7**.

**Table 3:** Parts required for the RemBot robot.

| Sl. No. | Part | Detail | Number of units/quantity |
| --- | --- | --- | --- |
| 1. | Microcontroller (Microchip ATmega 2560) | Controls the pumps and communicates to the secondary microcontroller (to control the temperature) | 1 unit |
| 2. | Secondary microcontroller (Microchip ATmega328p) | Controls the heating and cooling plates | 1 unit |
| 3. | 12V DC peristaltic pumps (7W) | Basic peristaltic pumps liquid dispersion and removal. Software controlled for precise buffer delivery. | 8 units |
| 4. | L298N driver board | Controls the peristaltic pumps and also provides a 5V supply. | 4 units |
| 5. | 2 channel 5v relay | Controls the dual heater plate from the secondary microcontroller. | 1 unit |
| 6. | 1 channel 5v relay | Controls when to heat the thermal plates. | 2 units |
| 7. | LCD2004 Module I2C Interface 5V | Displays the temperatures. | 1 unit |
| 8. | 2-inch TFT Screen (Driver: ST7789, 240x320 pixels) | The main display that shows the steps (with animations). | 1 unit |
| 9. | Colored wires male-female (50cms) | For connection. | 30 wires |
| 10. | Colored wires male-female (30cms) | For connection. | 40 wires |
| 11. | Solid core wires (red, black, green, yellow, blue) | For connections that require higher current (such as during heating) | 10 meters |
| 12. | Translucent high performance | Main tubing that connect the | 5 meters |
|  | Silicone Tubing, (ID - 1mm; OD - 2mm) | source, chamber, and sink. |  |
| 13. | Translucent high performance Silicone Tubing, (ID – 1.6mm; OD - 3mm) | Tubing to join the 1mm tubes | 1 meter |
| 14 | Translucent high performance Silicone Tubing (ID – 2 mm; OD - 3mm) | Tubing for microcomb outlet. | 10 cms |
| 15. | Y-connector (3.5mm) | To connect the tubes. | 10 units |
| 16. | Heating element (21x35 mm, 12V, 25W) | To heat the plates. | 2 units |
| 17. | Thermistor+10k ohm resistor | Measures the temperature of the heating element. | 3 units |
| 18. | USB cables | USB-A to USB-B, and USB-A to USB-C | 2 unit |
| 19. | Power supply (8A, 12V) | To provide power to the system | 2 units |
| 20. | PLA+ (grey: 1 spool and white: 1 spool) | For 3D printing. | 2 kgs |
| 21. | 3D printer. | BambuLab H2D | 1 unit |

**Table 4.** CAD parts for the RemBot robot (CAD files available at: https://github.com/tdblab/RAMFISH/tree/main/RemBot_STL).

| Sl. No. | Part | Description |
| --- | --- | --- |
| 1. | Main body | Main body of the system that holds the electronics and the pumps. |
| 2. | Front panel | Holds the display and buttons to control the system. |
| 3. | Top panel | Holds the probe modules with the buffer at the top of the system. |
| 4. | Back panel | Back plate of the system. |
| 5. | Chamber body | Main chamber that holds the confocal plate, confocal dishes, and the buffer exchanges. |
| 6. | Chamber body bottom plate | Bottom plate of the chamber body. |
| 7. | Confocal plate (136x96 mm; 125x85 mm; 125x80 mm) | Confocal microscope specific plate (designed differently for different microscope models) |
| 8. | Magnetic clip | Magnetic clip to hold the confocal dishes in the plate. |
| 9. | Microcomb adapter | Adapter to allow inlet and outlet to the confocal dishes |
| 10. | Microcomb plate | Plate to allow inlet and outlet to the confocal dishes (dual format) |
| 11. | Microcomb holder (free floating) | Dish holder for reactions that are carried out in free floating medium. |
| 12. | Microcomb holder (gel based) | Dish holder for reactions that are carried out in acrylamide gel. |
| 13. | Probe module plate 1 (Buffer plate) | Plate to hold the buffers. |
| 14. | Probe module plate 2 | Top lid of the buffer plate connected to the tubes. |
| 15. | Bottom cylinder support | Bottom cylindrical legs of the system. |

**Figure 5:**
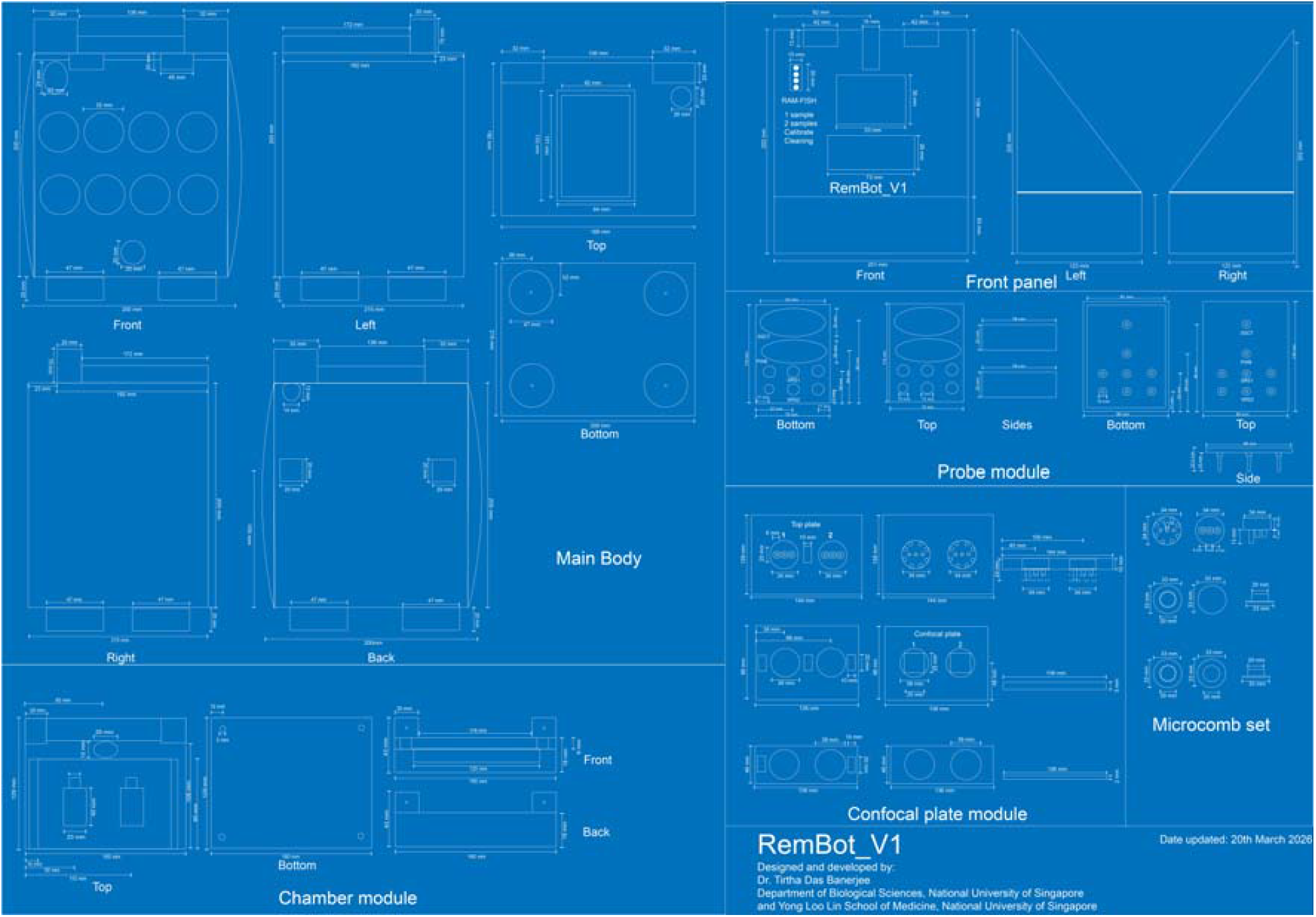
Architecture of the outer hardware of the RemBot robot. The components include the main body, front panel, probe module, confocal plate module, microcomb set, and chamber module.

**Figure 6:**
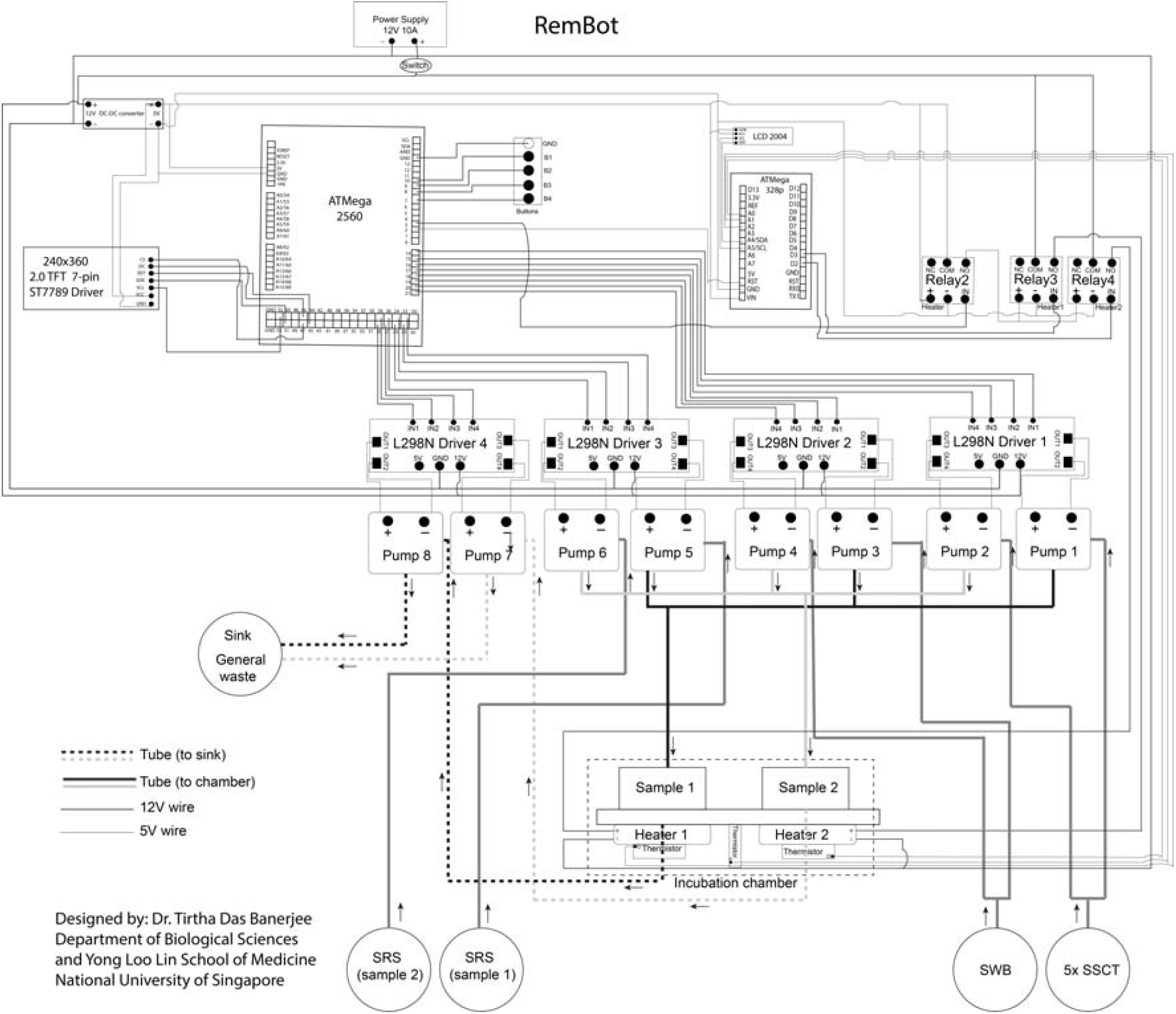
Circuit diagram of RemBot robot. The architecture integrates multiple pumps, sensors, and heating plates into an integrated system for carrying out the reactions.

**Figure 7:**
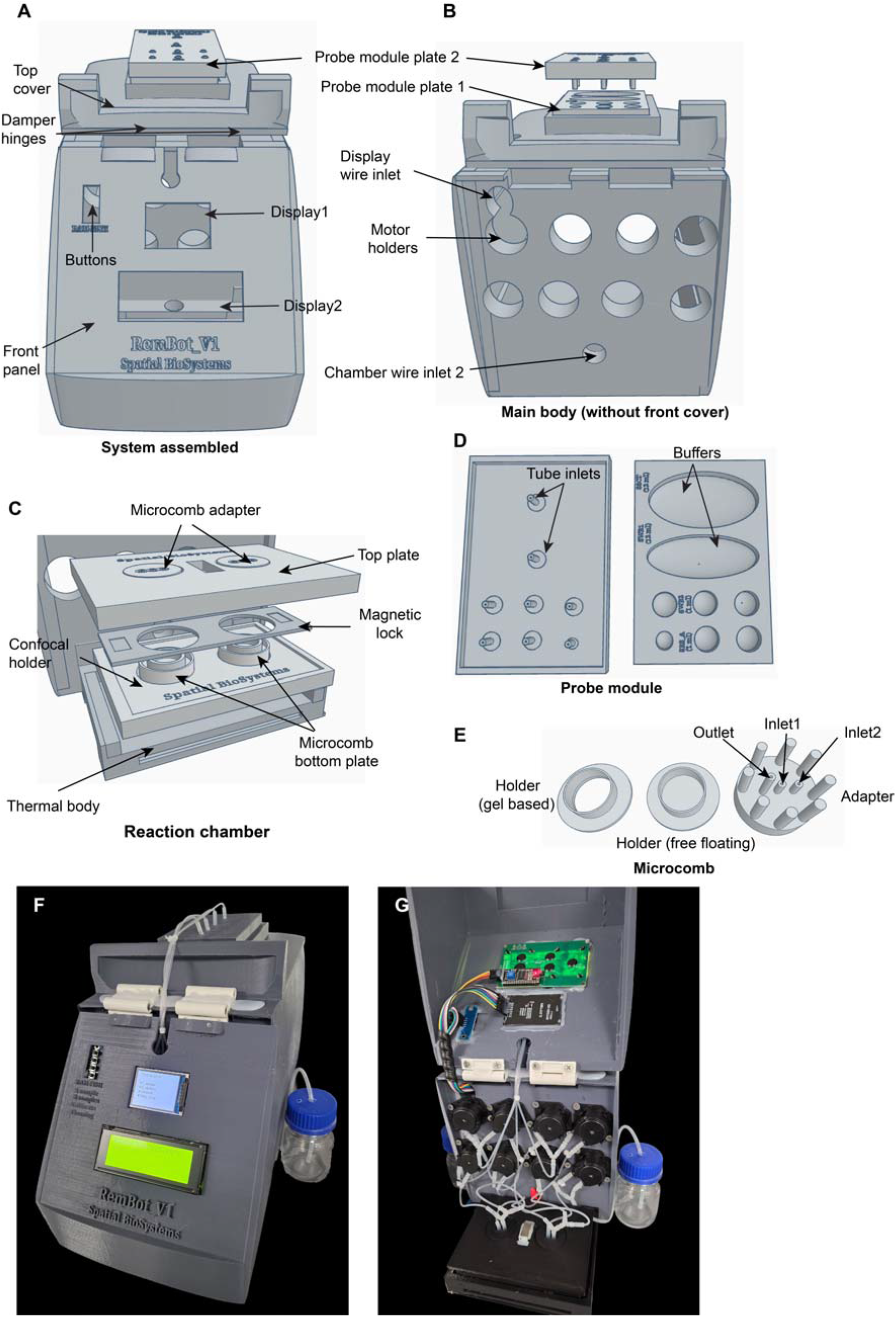
CAD design of the RemBot robot. **(A)** The main body of the robot. (**B**) Main body without the front cover showing the pump motor holders and the wire inlets. (**C**) Reaction chamber showing the microcomb top plate, magnetic lock, reaction surfaces, and the bottom plate. The thermal body holds the heating and cooling pads. (**D**) Probe module showing the plate to load the reaction buffers and the tube connectors from the top plate. (**E**) The microcomb module showing the bottom plate and top comb. An assembled RemBot-V1 robot with (**F**) closed front panel and (**G**) open front panel.

##### Step-by-step instruction for the RemBot robot assembly

Download all the STL files from the GitHub links provided in **Table 4** and 3D print them. For the assembly of all the STL files and the components in **Table 3**, follow the wiring architecture in **Figure 6** and the step-by-step instructions as mentioned below:

#### Main body

1. Solder wires to the peristaltic pumps and attach the pumps using screws to the main body.
2. Attach the pump wires to the L298N driver’s pins. Keep a note of the orientation of the connections. Two pumps can be connected to a single driver.
3. Connect the 12 V of the L298N drivers to a common 12V and the GND of the drivers to a common GND. The 12V common is connected to one side of the power switch; the other connection is to the power supply unit.
4. Connect the data pins from the L298N controllers to the IO pins of the Mega 2560 board (Pins: 3-6 and 14-44).
5. Connect the 12V common and GND to the input of the DC-DC voltage converter, adjust the variable resistor to match 5V using a multimeter as output. Connect the output 5V and GND to the common 5V and GND.
6. Install the 2-channel and 1-channel 5V relays using double sided tape in the main body and connect the 5V and GND from the common 5V and GND from the DC-DC converter. Connect the IN of the 1-channel 5V relay to pin 3 of the Mega 2560 board.

#### Front panel

7 Install the IPS and LCD screen with the attached female-to-male wire to the front panel using hot glue.
8 Connect the 5V and GND to the VCCs and GNDs of the microcontrollers, displays, and thermistors.
9 Connect the data pins of the IPS display to the IO of the Mega 2560 board (Pin: 49-59).
10 Connect the data pins of the LCD display to the I2C pins (A4 and A5) on the Nano 328p board.

#### Connecting the main body and the front panel

11 Attach the main body to the front panel using the damping hinge. The hinges can be connected using the self-tapping screws (4.2×13mm).
12 Attach the bottom cylindrical supports to the main body using the self-tapping screws (4.2×19mm).

#### Chamber

13 Attach the resistive heaters and the thermistors+10kΩ resistors assembly in the two slots provided in the chamber body.
14 Attach the chamber body bottom plate to the chamber body using self-tapping screws (4.2×13mm). Make sure the wires are accessible for later connection.
15 Attach the chamber body to the main body. Make sure the chamber body is horizontally aligned.
16 Connect 12V to the COM of the 2-channel 5V relays and NO channels to the two heaters 12V. Connect the GND from the heaters to the common GND. The IN pins from the relays are connected to the D2 and D3 pins, and the thermistor data pins to the A0 and A2 pins of the Nano 328p board.

#### Connecting the top, back, and main body

17 Connect the top panel to the main body using self-tapping screws (4.2×32mm) and the back panel to the main body using self-tapping screws (4.2×19mm). The USB cables are attached to both the Mega 2560 and Nano 328p boards before attaching the back panel.

#### Tubing assembly

18 Cut the tubes with 1.6mm ID into 1cm pieces and attach them to the outlets of the probe module plate 2. Connect the 1mm ID tubes to the 1.6mm tubes and join them to the inlets of the peristaltic pumps. Use Y-connectors with the 1.6mm ID tubes for the buffers that are shared with two pumps.
19 Connect the outlets from the pumps to the inlets of the microcomb adapters using 1mm ID tubes, 1.6mm ID tubes, and the Y-connectors.
20 Connect the outlets from the microcomb adapters to the outlet pumps (7 and 8) using 1mm ID tubes, 1.6mm ID tubes, and the Y-connectors to a 50 ml glass waste bottle.

#### Microcomb plate assembly

21 Attach fine nylon mesh (0.05 mm) to the output end of the microcomb using a 2 mm ID silicone tube (1mm height).
22 Attach the microcomb to the microcomb plate using hot glue.
23 Connect the inlets and outlets of the tubes to the microcomb inlets and outlets.
24 Add the confocal plate to the top of the chamber and install the microcomb plate.

### Uploading the control software

Download the codes for running single and dual samples for the multiplexer system from the GitHub link: https://github.com/tdblab/RAMFISH/tree/main/RemBot_code (rembot.ino and rembot_thermals.ino). Download and install Arduino IDE from the official site (https://support.arduino.cc/hc/en-us/articles/360019833020-Download-and-install-Arduino-IDE) on your computer. Install the necessary libraries(LCDWIKI_GUI.h, LCDWIKI_SPI.h, LiquidCrystal_I2C.h). Connect the USB-A ends of the USB cables to one of the PCs USB ports and the USB-C/USB-B ends to the boards. First, select the Mega 2560 board and upload the rembot.ino code. Next, select the Nano 328p board and upload the rembot_thermals.ino code.

## Results

In the present work, we tested the spatial expression patterns of five genes, *neurod1, gad2, nrp1a, oxt*, and *tph2. neurod1* (*neurogenic differentiation 1*) encodes a transcription factor critical to neuronal differentiation and widely expressed in the brain. Here, it is observed predominantly in the telencephalon, pineal gland, torus longitudinalis (medial border of Optic tectum), corpus cerebelli, and medulla oblongata tissues (**Figure 8A**). *gad2* (glutamic acid decarboxylase) encodes the GABA synthesis enzyme GAD65, facilitating production of the neurotransmitter. It also exhibits broad expression in the telencephalon, optic tectum, corpus cerebelli, and medulla oblongata (**Figure 8B**). *nrp1a* (*neuropilin 1a*) encodes a cell-surface receptor protein for semaphorins and vascular endothelial growth factor and plays a role in neuronal guidance. As seen here, *nrp1a* is expressed in the left dorsal habenula (**Figure 8C**) (Kuan et al., 2007), where it contributes to the asymmetric development of the habenula. *oxt* (*oxytocin/neurophysin I* prepropeptide) encodes the precursor protein required for the synthesis of oxytocin, and is known to be expressed in the preoptic hypothalamus (**Figure 8F**) (Nunes et al., 2021), as we observed. Finally, *tph2* (*tryptophan hydroxylase 2*) encodes for an enzyme critical to serotonin production, and is expressed in the pineal gland, dorsal thalamus, and dorsal raphe (**Figure 8G**). The observed signals for each gene assayed match closely with published data in larval zebrafish (Shainer et al., 2023), and indicate the applicability of the technique to both surface level and deep-seated targets in whole mount tissues.

**Figure 8.**
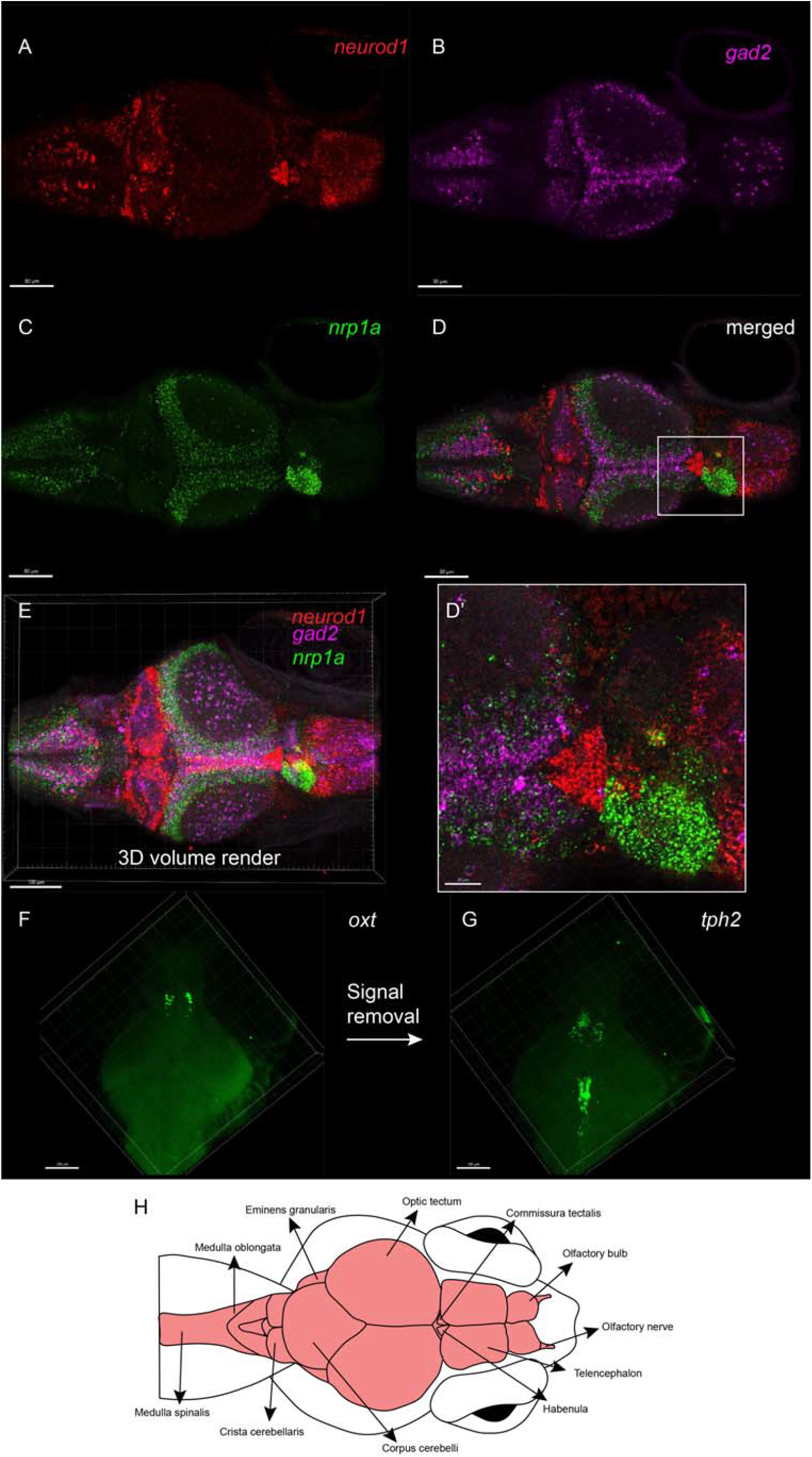
Expression of *neurod1, gad2, nrp1a, oxt*, and *tph2* in 14 dpf old zebrafish larvae carried out using RAMFISH protocol using the automation module. (**A-D**) Single slice (∼5 μm) of a 14 dpf zebrafish larva expressing *neurod1, gad2*, and *nrp1a* and (**E**) a 3D volume render (∼200 μm thick). Expression of (**F**) *oxt* in a 3D volume rendered zebrafish (∼200 μm thick) followed by signal removal and stained against (**G**) *tph2*. **(H)** Illustration of a zebrafish brain.

A detailed and comprehensive list of genes tested are available at Banerjee et al., 2024.

## Discussion

The automation platform described here provides a flexible and accessible solution for executing the RAM-FISH workflow (Banerjee et al., 2024) that requires repeated reagent exchanges, controlled incubation conditions, and precise timing. By combining programmable fluid delivery with integrated thermal control to accelerate the reaction time and movement of DNA probes (**Figure S1-S3**), our robots enable reliable execution of cyclic experimental protocols directly in microscopy dishes. The modular architecture allows the robots to be adapted for various needs: free-floating and gel embedded tissue preparations, for example, while widely available components and open-source control software makes them practical for implementation in research laboratories without engineering infrastructure or expertise. The approach and the design prioritize adaptability and cost-efficiency, in contrast to commercial systems optimized for specific consumables or proprietary workflows.

While the current dual-robot system was validated using the RAM-FISH workflow, the underlying architecture is readily compatible with a range of iterative labeling protocols that involve sequential reagent delivery and washing steps. The platform currently has a few limitations, including optimization of fluidic precision for very small reaction volumes (50 μl or less) and expansion of temperature control capabilities for additional reaction conditions. Future improvements will include integration with microscope control software via direct communication with the microcontroller, expanded multi-channel fluidic routing by adding additional pumps and valves for multi-round staining, and further miniaturization of the reaction chamber for direct attachment to a confocal system. Overall, this system provides a practical open-source framework for automating mRNA profiling and lowers technical barriers to multi-cycle *in-situ* hybridization experiments.

## Method

14 dpf zebrafish (Danio rerio) were housed, bred, and reared at the Zebrafish Facility (ZFF, Institute of Molecular and Cell Biology, A*STAR) in groups of 20–30 in 3 L tanks at 28°C. Fish were kept under a 14:10 light-dark cycle and fed as per the standard operating procedures of ZFF.. Spatial multiplexing on the genes described was carried out based on RAMFISH based automation protocol described in Banerjee et al., 2024. Briefly, zebrafish larvae collected were 1) fixed in 4% formaldehyde, 2) washed with 1x PBST, 3) permeabilized using SDS/ammonium lauryl sulfate based permeabilization solution, 4) washed with 1x SSCT, and 5) finally transferred in 20% Ethylene carbonate based or 30% probe hybridization buffer. Afterwards, buffer were loaded into the wells of the plate (probe_module_1) and the larvae were transferred to wells (microcomb_holder_free_floating) of the Multiplexer system, and the script for the standard protocol was executed (8 hours in total). Imaging was carried out in Olympus FV3000 and Zeiss LSM700 confocal microscopes. After imaging, signals were removed using a DNase I-based buffer described in Banerjee et al., 2024 and the signal localization reaction was repeated.

## Authors contribution

TDB: Conceptualization, hardware design, programming, simulations, methodology, writing - original draft, writing -review and editing. JR: Methodology, writing - review and editing. ASM: Supervision, writing - review and editing, funding. AM: Supervision, funding, writing - review, and editing.

## Acknowledgement

We thank DBS-CBIS confocal facility and Tong Yan, for access to the Olympus FV3000 confocal microscope, and Tan Lu Wee for lab management.

## Data availability

The software and CAD designs in the present manuscript can be found in the GitHub link: https://github.com/tdblab/RAMFISH.

## Associated Article

RAMFISH workflow and results are available at https://www.biorxiv.org/content/10.1101/2024.12.06.627193v2

