## Supplementary File 1 for "Modular fluidic automation platform with integrated thermal control for multi-step molecular imaging workflows"

Tirtha Das Banerjee^1,3,*^, Joshua Raine^2^, Ajay S. Mathuru^2,3,4,5,*^, Antónia Monteiro^1,*^

### Affiliations

^1^Department of Biological Sciences, National University of Singapore 117557, Singapore.

^2^ Department of Physiology, Yong Loo Lin School of Medicine, National University of Singapore, Singapore.

^3^ Institute for Digital Medicine (WisDM), Yong Loo Lin School of Medicine, National University of Singapore, Singapore.

^4^ N.1 Institute for Health, National University of Singapore, Singapore.

^5^ Healthy Longevity TRP, Yong Loo Lin School of Medicine, National University of Singapore, Singapore.

*Authors for correspondence:

**Mean Squared Displacement (MSD) and DNA particle simulation**

In this supplementary file we calculated the difference in the movement of the DNA particles in three different thermal fields at 25°C, 37°C, and 45°C in water medium; and generated simulation using the known thermal diffusivity and diffusion values of small DNA oligos (Duhr and Braun, 2006).

**MSD calculations**

The mean squared displacement (MSD) quantifies the average squared distance particles have traveled from their initial positions. This is a useful parameter that can help us identify the speed at which the particles (DNA) are moving over a period of time inside the chamber in each scenario.

Calculation of Diffusion at different temperatures

Diffusion of small molecules typically scales with viscosity and temperature via the Stokes-Einstein relation (Einstein, 1905; Stokes, 2010)	.

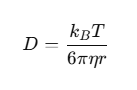

**Table S1: Parameters for Stokes-Einstein equation and their units.**

| Symbol | Description | Units |
| --- | --- | --- |
| D | Diffusion coefficient | m²/s |
| k_B_ | Boltzmann constant | 1.380649×10^−23^ J/K |
| T | Absolute temperature | K |
| η | Dynamic viscosity | Pa·s |
| r | Hydrodynamic radius of the particle | m |

Base diffusion (D_base_) = 0.85x10^-6^ cm^2^/s at 25°C.

This value is assumed based on typical diffusion coefficients for small DNA oligos (HCR probes are 45 bps and secondary probes are 72 bps (in hairpin structure)) in aqueous solution at room temperature. DNA fragments of ~30–50 base pairs have diffusion coefficients in the range of 7x10^-7^ to 1x10^-6^ cm^2^/s at 25°C. (Duhr and Braun, 2006).

Diffusion coefficients at different temperatures were calculated using Stokes-Einstein relation (Einstein, 1905; Stokes, 2010).

D_1_/D_2_ = T_1_/T_2_ x η_2_/ η_1_

η _25_ = Dynamic viscosity of water at 25°C: 0.89 mPa·s

η _37_ = Dynamic viscosity of water at 37°C: ~0.691 mPa·s

η _45_ = Dynamic viscosity of water at 45°C: ~0.596 mPa·s

Diffusion at 37°C

D_37_ = D_25_ x (37+273.15)/(25+273.15) x η _25_/ η _37_

= 0.85x10^-6^ x 1.04 x 1.28 = 1.14 x 10^-6^ cm^2^/s

Diffusion at 45°C

D_45_ = D_25_ x (45+273.15)/(25+273.15) x η _25_/ η _45_

= 0.85x10^-6^ x 1.07 x 1.49 = 1.36 x 10^-6^ cm^2^/s

**Mean square displacement**

In water

Case 1: Heat at RT =25°C

Time (t) = 200 sec

Temperature: constant 25°C

Diffusion at temperature 25°C

D_25_ =0.97×10^−6^ cm^2^/s

MSD_diff (25)_ = 4D_25_t

= 4×0.85×10^-6^ ×200 = 6.88×10^−4^ cm^2^

Case 2: Heat at 37°C

Time (t) = 200 sec

Temperature: constant 37°C

Diffusion at temperature 37°C

D_37_ =1.14×10^−6^ cm^2^/s

MSD_dif(37)f_ = 4D_37_t

= 4×1.14×10^−6^×200 = 9.12×10^−4^ cm^2^

Case 3: Heat at 45°C

Time (t) = 200 sec

Temperature: constant 37°C

Diffusion at temperature 37°C

D_37_ =1.36×10^−6^ cm^2^/s

MSD_dif(37)f_ = 4D_37_t

= 4×1.36×10^−6^×200 = 10.88×10^−4^ cm^2^

**Simulations**

We simulated the thermally driven movement of DNA particles in a 2D microchamber (3.5 × 3.5 cm), incorporating both heat diffusion and thermophoretic particle drift. In all cases, the temperature field evolved using explicit finite difference solutions of the heat diffusion equation, while DNA particles underwent Brownian motion with temperature-dependent diffusivity, combined with lateral drift driven by local temperature gradients.

The thermal diffusivity used in simulations was 0.00143 cm²/s, based on empirical measurements for water as below.

Viscosity (*µ*) at 25°C = 1 mPa.s =0.001 Pa.s

density (*ρ*) at 25°C = 1 kg/m^3^

Kinematic viscosity (*ν*) = *µ/* *ρ* = 0.001/1000 = 1 x 10^-6^ m^2^/s

Thermal conductivity k = 0.6 W/(m.K)

Specific heat Cp = 4182 J/(kg.K)

Thermal diffusivity (α) = *k*/ ρC_p_ = 0.6/(1000x4182) = 1.43 x10 ^-7^ m^2^/s

**Simulation1: Uniform heat in the reaction chamber (25°C)**

Initial Temperature: 25°C (uniform throughout chamber)

Boundary Conditions: Constant 25°C on all sides

Particles: 200 DNA probe-sized particles with temperature-dependent diffusion.

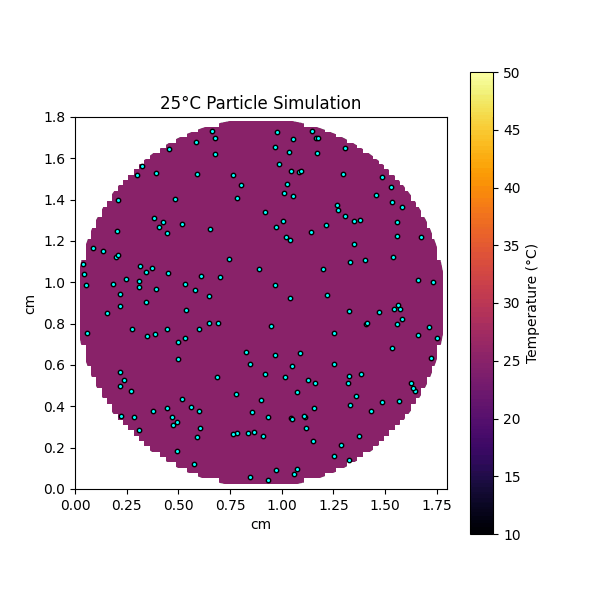

**Figure S1. Movement of particles at 25°C** **thermal field due to Brownian movement.**

**Simulation2: Uniform heat in the reaction chamber (37°C)**

Initial Temperature: 37°C (uniform throughout chamber)

Boundary Conditions: Constant 37°C on all sides

Particles: 200 DNA probe-sized particles with temperature-dependent diffusion.

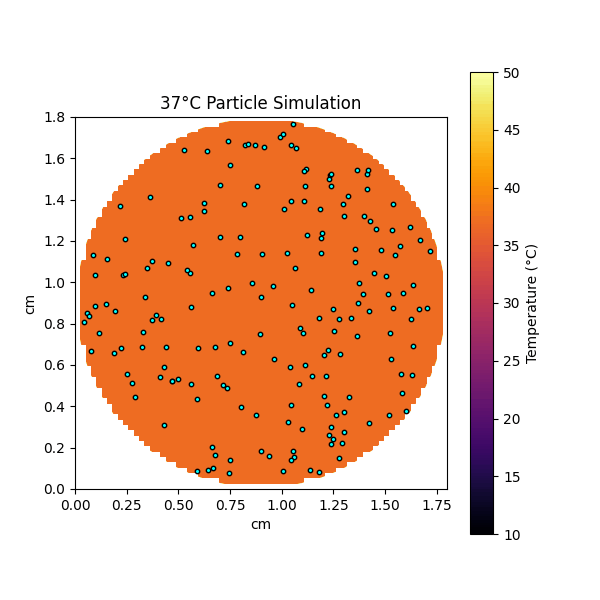

**Figure S2. Movement of particles at 37°C** **thermal field due to Brownian movement.**

**Simulation3: Uniform heat in the reaction chamber (45°C)**

Initial Temperature: 45°C (uniform throughout chamber)

Boundary Conditions: Constant 45°C on all sides

Particles: 200 DNA probe-sized particles with temperature-dependent diffusion.

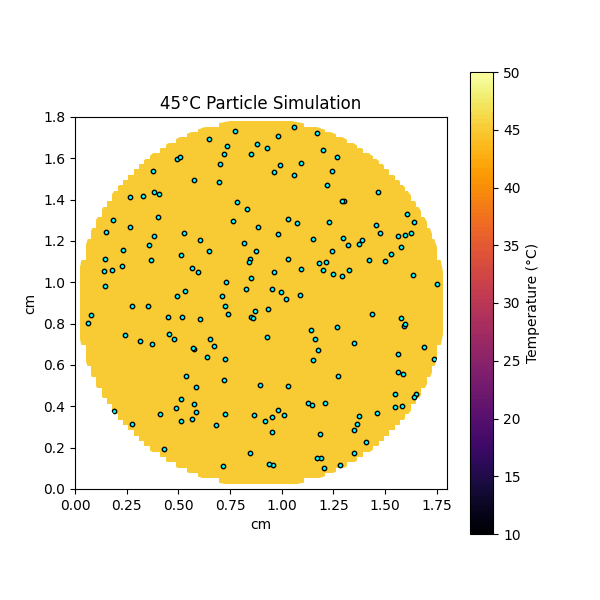

**Figure S3. Movement of particles at 45°C** **thermal field due to Brownian movement.**
